# DMB labelling for detection and analysis of capsular polysaccharides

**DOI:** 10.1101/2025.01.29.635457

**Authors:** Christoph Rutschmann, Kateryna Vershynina, Corina Mathew, Lucas Piëch, Małgorzata Sulewska, Philipp E Schilling, Nicolas Wenner, Louise Larsson, Anouk Bertola, Elisa Cappio Barazzone, Ioana Domocos, Carine Roese Mores, Jackie Raisse Ndem-Galbert, Yagmur Turgay, Raffael Schumann, Shirin de Viragh, Hayley Lavender, Chris Tang, Médéric Diard, Shinichi Sunagawa, Michael Wetter, Timm Fiebig, Emma Slack, Timothy G Keys

## Abstract

Bacterial capsules are major virulence factors enabling systemic infection by undermining innate and adaptive immunity. Capsular polysaccharides are also the antigen in some of the most successful antibacterial vaccines – including the pneumococcal, neisserial and Hib conjugate vaccines. However, it remains exceptionally challenging to study capsules, primarily due to their high chemical diversity and the limited methods available for their detection and analysis. We describe a robust biochemical method for detection and analysis of ABC transporter-dependent capsular polysaccharides, a major class of capsules that are associated with extraintestinal pathogenic *Escherichia coli* (ExPEC). The method involves release and fluorescent tagging of polysaccharides from the cell surface. Anion exchange chromatography of labelled samples reveals the presence, relative abundance, and length-distribution of these diverse polysaccharide antigens. The method provides a modern approach to detecting the capsule phenotype, bridging a critical gap left by the decline of serotyping assays. It will enhance our understanding of fundamental capsule biology and advance the development of capsule-targeting vaccines.

## Main

Bacterial capsular polysaccharides are known for their roles in virulence [1]–[4], bacteriophage evasion and predation [5]–[8], survival in harsh environments [9], [10], and as antigens in glycoconjugate vaccines [11]–[13]. The study of these important molecules is hindered by a lack of methods for their facile detection and analysis.

*Extraintestinal pathogenic Escherichia coli* (ExPEC) is a leading cause of antimicrobial resistance associated deaths and has been identified by the WHO as a top-priority pathogen for vaccine R&D [14], [15]. Polysaccharide capsules play an essential role in invasive ExPEC infections by mediating resistance to complement and phagocytic killing [16]. This function is supported by the inherent poor immunogenicity of capsules, such that even invasive infections fail to induce considerable anti-capsule antibody responses [17], [18]. On the other hand, we can exploit the prominent location of these polysaccharides on the ExPEC surface. Capsule targeting antibodies were shown to protect against ExPEC infection more than 40 years ago [19], [20].

Despite the biomedical importance of capsules, there is no method available for their routine detection or analysis. Current methods are limited to a serotyping assay known as known as counter current immunoelectrophoresis [21]. This method was central to establishing our initial understanding and catalogue of K antigens [22]. However, the study of *E. coli* capsule serotypes declined together with the field of serology after the rise of sequence-based methods for identifying and tracking isolates (e.g. MLST) [23]. Today, the antisera required for capsule serotyping have only restricted commercial availability, and we are only aware of a single company offering K-serotyping services for *E. coli*. As a result, we have little modern data on the association of capsules with *E. coli* lineages or pathotypes [24]–[26]. Furthermore, despite genetic evidence for new and emerging capsule types, no new K antigens have been identified since 1977 [25]–[28].

We aimed to remove this barrier by developing a simple biochemical method for detecting transporter-dependent capsular polysaccharides, that are associated with the ExPEC pathovar. Transporter-dependent (also known as group 2 and 3) polysaccharides have negatively charged repeat units and a common glycolipid anchor composed of a β-linked oligosaccharide of the α-keto acid sugar, Kdo (3-deoxy-D-*manno*-oct-2-ulosonic acid) attached to a diacylglycerol phosphate lipid [29]–[31]. Kdo linkages are subject to hydrolysis under mild acidic conditions, providing a possible mechanism for releasing these polysaccharides from the bacterial surface [32]. Furthermore, Kdo can be fluorescently labelled with the α-keto acid-specific reagent, DMB (1,2-diamino-4,5-methylenedioxybenzene), a bright fluorophore that has transformed the analysis of polysialic acids, enabling sensitive enzymatic assays, and the detection of polysaccharide directly from complex tissue samples [33]–[37]. Given these points, we reasoned that it might be possible to release and DMB-label capsular polysaccharides to facilitate their detection and analysis directly from complex whole-cell samples (Fig. 1a).

**Figure 1.**
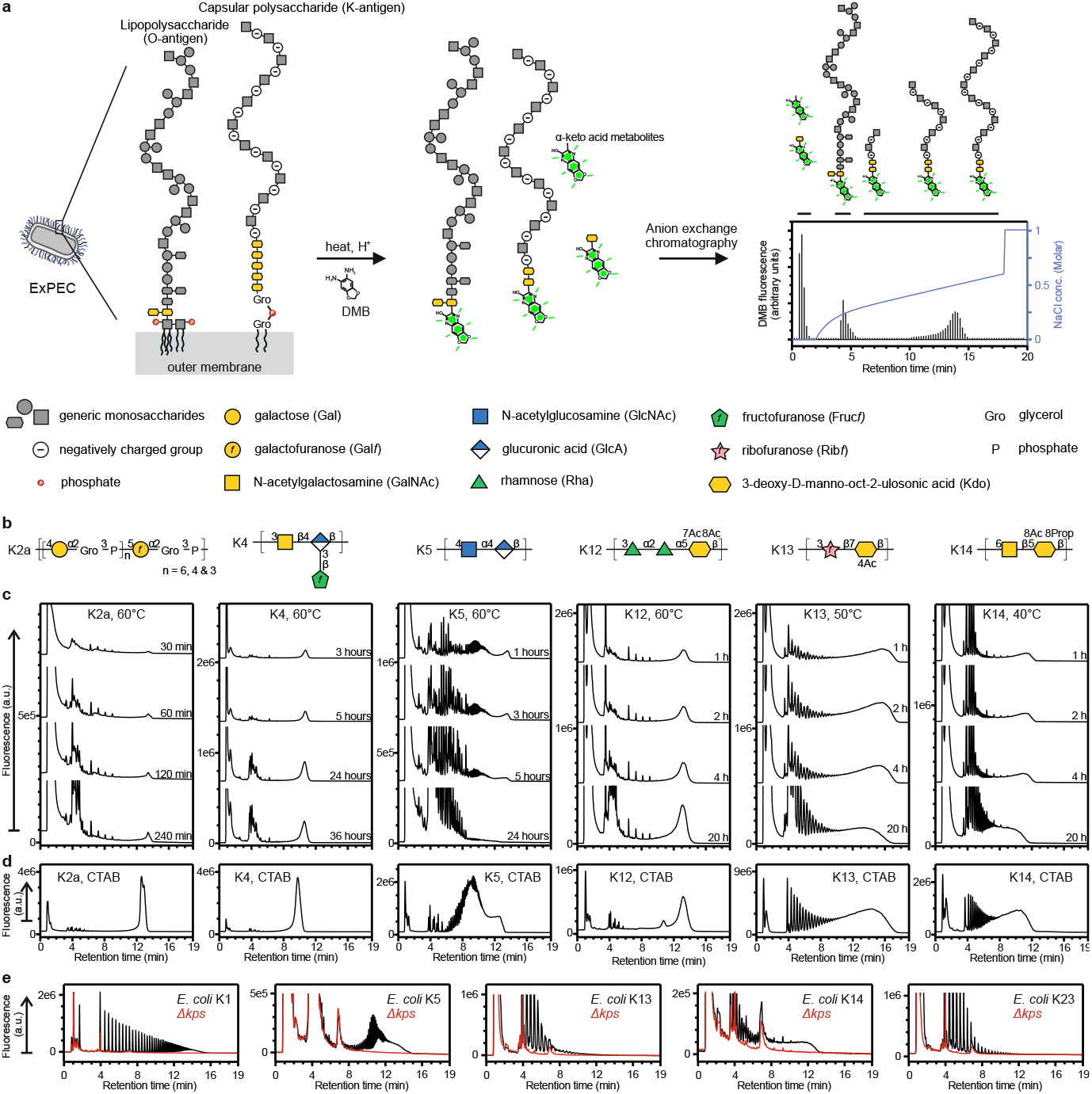
DMB labelling of capsular polysaccharides. (**a**) Schematic overview of DMB labelling of whole-cell samples. Lipopolysaccharide and capsular polysaccharide are ExPEC’s major surface antigens. These polysaccharides can be released from the outer membrane by hydrolysis of Kdo-linkages under mild acidic conditions. In the presence of DMB, the reducing-end of Kdo residues and α-keto acid metabolites are fluorescently labelled. When separated by anion exchange chromatography, neutral and weakly charged species, including O antigen, run at low retention times. The polyanionic capsular polysaccharides are separated according to charge and length with long chain species running at high retention time. (**b**) K antigen repeat unit structures in symbol nomenclature [89]. (**c**) Whole bacterial cells were incubated in 50 mM TFA at the indicated temperature. Samples were taken over time, supernatants were DMB labelled, then analyzed by anion exchange chromatography with online monitoring of DMB fluorescence. (**d**) Capsular polysaccharides were purified from the reference strains, DMB labelled, then analyzed by anion exchange chromatography. (**e**) The capsule biosynthesis locus was deleted (*Δkps*) from *E. coli* with the K1, K5, K13, K14, and K23 antigens. The wild-type and corresponding *Δkps* strains were cultivated in AUM, cell pellets were DMB labelled, and supernatants were analyzed by anion exchange chromatography.

## Results

To test hydrolysis and DMB labelling we chose a structurally diverse set of K antigens (Fig. 1b). These included a glycerol-phosphate-containing polysaccharide (K2a), two polysaccharides with glycosaminoglycan backbones (K4 and K5), and three polysaccharides with different Kdo-containing repeat units (K12, K13, and K14). We explored the time and temperature required to release each polysaccharide from whole cells in mild acidic solution. Material released into the supernatant was DMB labelled, then separated by anion exchange chromatography with online fluorescence detection (Fig. 1c). In each case we observed DMB-labelled species at high retention time (>10 minutes), consistent with our expectations for polyanionic capsular polysaccharides. The putative capsular polysaccharides exhibited different peak profiles and different stability to the hydrolysis conditions.

We pursued biochemical and genetic strategies to verify the identity of the strongly retained species. DMB labelling of purified capsular polysaccharides yielded fluorescence profiles in excellent agreement with the strongly retained species observed from cell hydrolysates (Fig. 1d). Furthermore, deletion of the *kps* locus, including the full set of genes responsible for capsule biosynthesis and export, resulted in a corresponding ablation of the strongly retained species (Fig. 1e). These results demonstrate that capsular polysaccharides can be released from whole cells by mild acid hydrolysis, fluorescently labelled with DMB, and specifically detected as species with high retention time in anion exchange chromatography.

To assess whether the method is broadly applicable to transporter-dependent capsules, we labelled a collection of *E. coli* K antigen reference strains. We observed putative capsular polysaccharide from 30 of 31 strains that are reported to produce a transporter-dependent capsule (Supplementary Data Set 1). No signals resembling capsular polysaccharide were observed from unencapsulated negative control strains. The results demonstrate that DMB labelling is broadly, though not universally, applicable to the chemically diverse family of transporter-dependent capsular polysaccharides.

We did not observe a DMB signal at high retention time from the K3 reference strain. The K3 and K98 polysaccharides have an identical neutrally charged poly-rhamnose backbone. These polysaccharides acquire negative charge by modification with hexulosonic acid and glucuronic acid side-branches, respectively. The unique linkage attaching the hexulosonic acid side-branch to the K3 polysaccharide was previously observed to be a “very labile linkage” [38]. Therefore, hydrolysis of this bond, and loss of negative charges during labelling, is the most likely reason for failure to observe the DMB-labelled K3 polysaccharide at high retention times. This highlights a specific limitation of DMB labelling for the analysis of polysaccharides with rare bonds that are more labile than glycosidic bonds of Kdo [32].

To explore the specificity of the DMB labelling strategy, we analyzed *E. coli* reference strains that produce Wzx/Wzy-dependent K antigens (also known as group 1 and 4 capsules). From 40 Wzx/Wzy-dependent K-types, two, K9 and K55, were observed to give a transporter-dependent-like signal (Supplementary Data Set 1). The K9 polysaccharide is subject to hydrolysis and DMB labelling due to the presence of an α-keto acid sugar, sialic acid, in the repeat unit. The nature of the labelled species from the K55 strain is not clear, however it may be that the K55 antigen is efficiently attached to lipid-A, in a form known as K_LPS_ [39]. Notably, whole genome sequencing of the K9 and K55 strains confirmed the absence of genes associated with transporter-dependent capsules (*kpsCS*). The results indicate that DMB labelling results cannot be used in isolation to conclude on the presence of a transporter-dependent capsule. Combining the results with genome sequence information will be required to constrain the types of surface antigen that may be produced by a strain. Furthermore, the results highlight the potential for detection and discovery of further α-keto acid-containing bacterial molecules.

DMB labelling is robust to variations in the fine structure of the polysaccharide (e.g. variable levels of acetylation) and other properties of the bacterial strain (such as restriction modification systems or bacterial immunity). It should therefore provide a more reliable method for detection of capsular polysaccharides than serotyping assays. Especially in the case of K1 and K5, where the capsule phenotype is defined by capsule-specific bacteriophages that may be subject to capsule-independent resistance mechanisms [1], [7], [8], [40], [41]. To test this proposition, we compared phage-based serotyping and DMB labelling for five clinical isolates where genome sequencing revealed a K5-like *kps* locus (Fig. 2a). The K5 reference strain and the K5 commensal *E. coli* Nissle 1917 were included as positive controls. Interestingly, while only three of five clinical isolates were susceptible to the serotyping phage (Fig. 2b, Supplementary Information Fig. S1), DMB labelling revealed that four strains produced a capsular polysaccharide (Fig. 2c). We controlled the structure of the DMB-labelled polysaccharides by testing their susceptibility to digestion with a recombinant, K5-specific, bacteriophage tailspike lyase [42]. Polysaccharide from all strains was susceptible to K5-lyase digestion (Fig. 2c). We conclude that LU071 is a K5 strain, that gives a false-negative serotyping result due to a phage resistance mechanism that is independent of the capsule structure.

**Figure 2.**
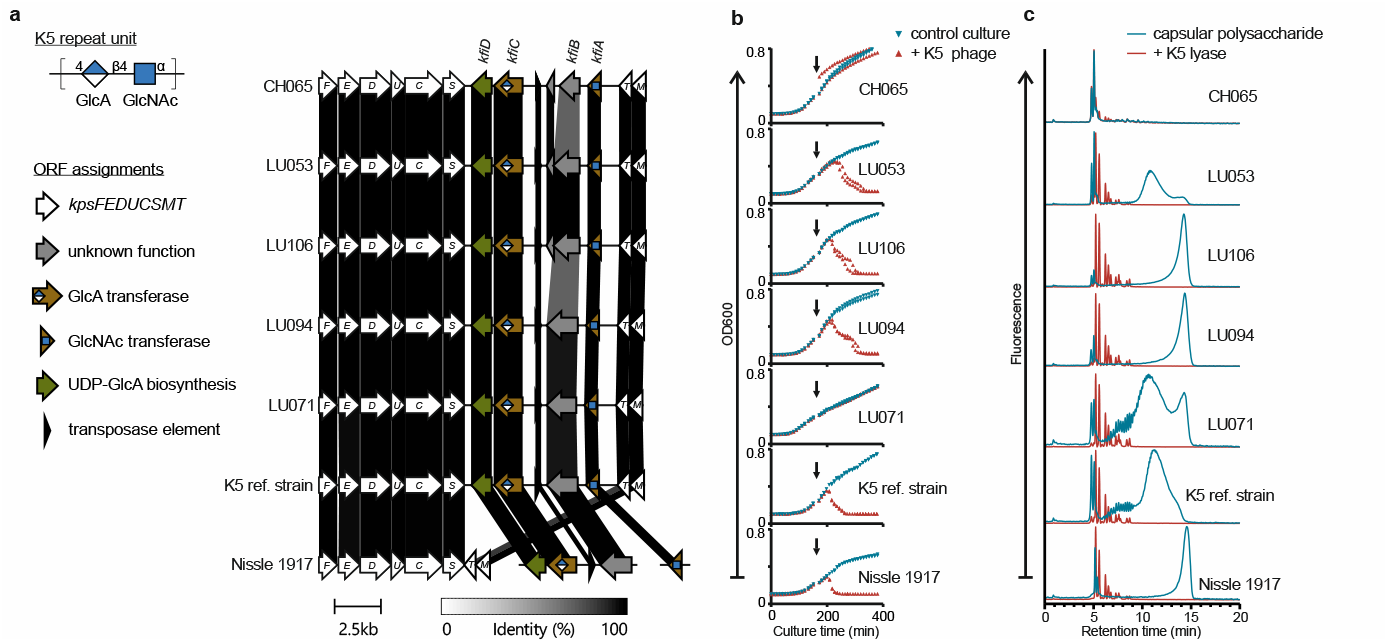
Serotyping and DMB labelling analysis of putative K5 clinical isolates. (**a**) Schematic comparison of capsule gene clusters from each strain. The *kps* locus was extracted from whole genome sequence data by fastKaptive [25] and compared to the K5 reference strain and *E. coli* Nissle 1917 with CLINKER [88]. (**b**) Each strain was tested for susceptibility to the K5 typing bacteriophage in liquid culture and using the cross-brush method (**Fig. S1**). Four independent cultures of each strain were grown in LB media at 37°C. K5 bacteriophage was added to two of the cultures at mid-log phase (black arrow). OD_600_ was measured every 10 minutes. All data points are shown. (**c**) Capsular polysaccharide was hydrolysed from each strain, separated from metabolites by ultrafiltration, DMB labelled, and analyzed by anion exchange chromatography. The K5 structure of each polysaccharide was verified by testing for susceptibility to depolymerization by a recombinant K5 lyase [42]. Reaction products were evaluated by anion exchange chromatography.

There is evidence for complex transcriptional-level regulation of capsule expression in pathogenesis of urinary tract infections and meningitis [43]–[49]. However, studies of capsule expression are generally limited to K1 and K5 capsules because molecular tools such as capsule-specific antibodies or engineered bacteriophage tailspike proteins are available to detect these polysaccharides [22], [50], [51], whereas such reagents are missing for basically all other serogroups. We are aware of a single experiment that compared expression of diverse K antigens under different cultivation conditions. This influential experiment, published in 1984 by Fritz and Ida Ørskov [52], used counter current immunoelectrophoresis to assess capsule expression after cultivation at 18°C or 37°C, and led to the paradigm that group 2 capsules are temperature-regulated while group 3 are expressed irrespective of cultivation temperature [53], [54]. We revisited this experiment by growing strains assigned to group 2 or 3 at temperatures from 16°C to 37°C and measuring capsular polysaccharide by DMB labelling (Fig. 3). The quantitative nature of DMB labelling reveals a more nuanced picture of capsule expression. We observe a spectrum of behaviours from relatively uniform expression at different temperatures to capsules that are only observed after cultivation at 37°C. A limitation of this experiment is that we have only assessed cell-associated capsular polysaccharide; any material sloughed into the media is lost to this analysis. We anticipate that DMB labelling will greatly facilitate future studies of capsule regulation in response to diverse environmental stimuli.

**Figure 3.**
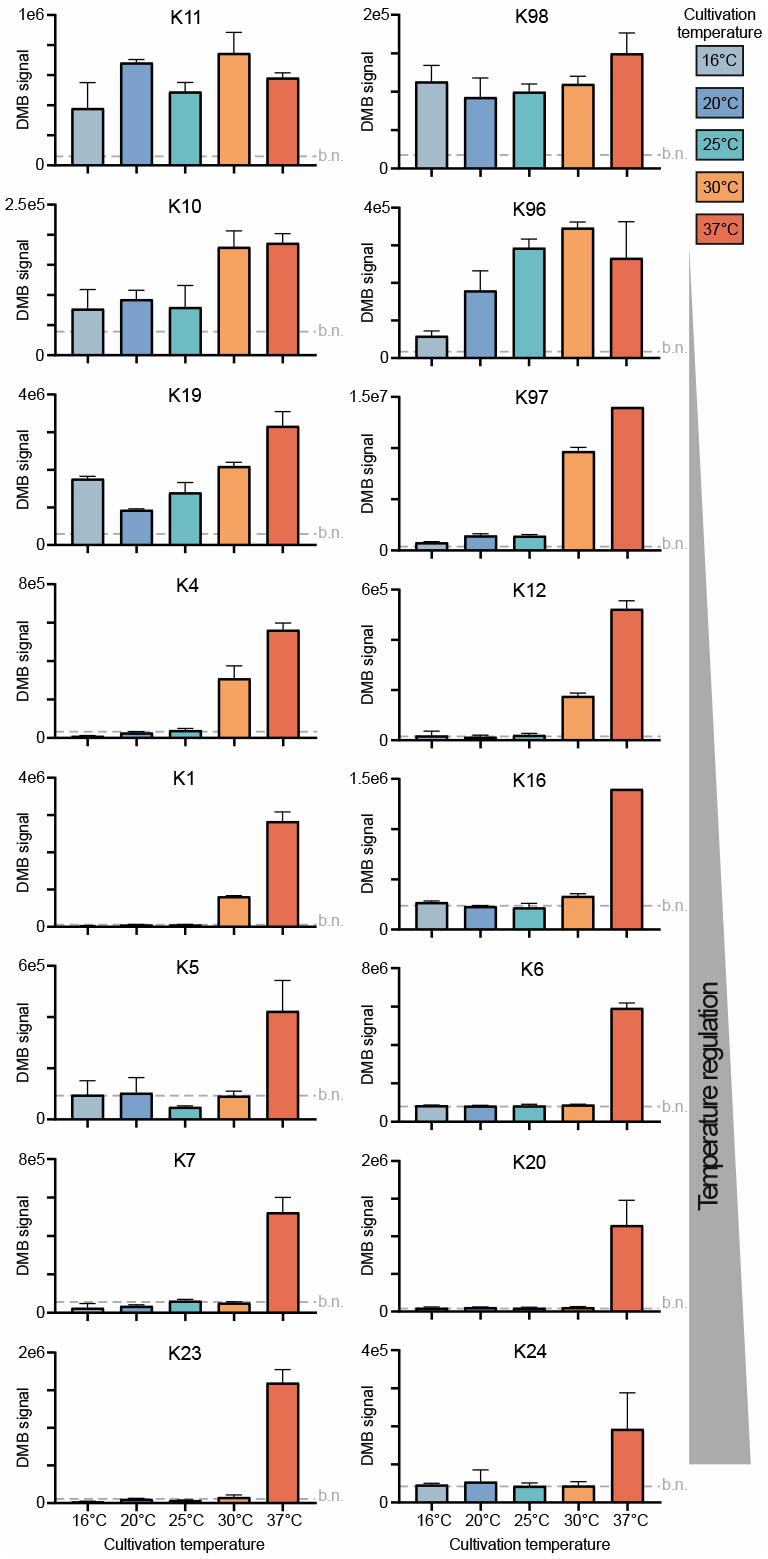
Temperature dependence of capsule expression. The K antigen reference strains were cultivated to stationary phase in TB media at the indicated temperatures. Cells were harvested, washed, then assessed for capsule by DMB labelling and anion exchange chromatography. The amount of capsule was estimated as the area under the curve (DMB signal) in a defined retention time range. The mean average and standard deviation of three cultures are plotted. The approximate level of background noise (b.n.) is indicated on each plot.

## Discussion

For most of the 20^th^ century, *E. coli* strains were identified by their major surface structures, the serologically defined O, H, and K antigens. However, the practice of serology declined with the advent of molecular methods for strain identification, like MLST [23]. This decline has been partially compensated by genotyping algorithms for O and H [55]–[57], but not for K antigens. As a result, the abundance, distribution, and disease-associations of O and H antigens is well studied [58]–[63], whereas similar information on most K antigens is lacking [26]. Furthermore, the repertoire of K antigens has not been updated since 1977 [28], despite evidence of uncatalogued capsule diversity [25], [27], [64]. Due to the lack of methods for detecting capsule expression, ExPEC vaccine studies, where capsule is acutely relevant, rely on the presence of capsule biosynthesis genes and observations of “O antigen masking” to infer capsule expression [65], [66].

To facilitate and reinvigorate the study of ExPEC capsular polysaccharides, we have demonstrated that DMB labelling combined with anion exchange chromatography constitutes a robust biochemical method for sensitive and quantitative detection of transporter-dependent capsular polysaccharides. The method is applicable to a variety of samples including whole cells, crude extracts, or purified polysaccharides, and it rapidly provides rich information including relative abundance and chain-length distribution of polysaccharides using standard HPLC equipment.

A limitation of DMB labelling is that the mild-acidic conditions result in partial loss of chain length information in polysaccharides that include labile α-keto acid sugars (e.g. sialic acid and Kdo) in the repeat unit (e.g. K1, K6, and K13). This can be largely mitigated by optimizing the hydrolysis conditions for the polysaccharide of interest. Nevertheless, as observed for K3, polysaccharides with rare linkages that are more labile than those of Kdo may be destroyed by the sample preparation process.

The primary advantage of DMB labelling over available methods is the ability to detect and quantify polysaccharides from a small sample of whole cells. Polyacrylamide gel electrophoresis (PAGE) with Alcian blue-silver staining and HPLC separations with UV detection or pulsed amperometric detection are excellent methods for analysis of pure polysaccharide samples [67]–[77]. However, purifying capsular polysaccharide is non-trivial and time-consuming, making PAGE and HPLC impractical for routine or high-throughput screening of strains for capsule production. Furthermore, PAGE and HPLC cannot rival the nanogram sensitivity that is achieved with the DMB label [34], [35].

DMB labelling does not allow for identification of specific K antigens and therefore does not serve as a tool for serotyping. However, for K-types with a published *kps* locus sequence [78], [79], genotyping can assign a K-type and the capsule phenotype can be confirmed by DMB labelling.

Furthermore, DMB labelling has several advantages compared to classical antisera-based serological assays: (i) it can be used to identify novel or emerging K antigens that do not have established antisera, (ii) it provides information on quantity and length of polysaccharides produced by any given strain, and (iii) it is not sensitive to minor structural variation of the polysaccharide, e.g. acetylation [22].

DMB labelling has proven useful for a wide variety of tasks from assessing encapsulation of clinical isolates to tracking polysaccharide through purification procedures and for verifying recombinant expression of capsular polysaccharides. We anticipate that DMB labelling will serve as a pillar of future research into capsule biology It will advance ExPEC vaccine development and complement studies of K antigen genetics to expand the catalogue of transporter-dependent capsules produced by *E. coli*.

## Methods

### Bacterial strains and cultivation conditions

All strains used or developed in this study are listed in Supplementary Table 1. Unless otherwise indicated, strains were cultivated in lysogeny broth (LB) at 37°C without antibiotic selection, liquid cultures were shaken at 180 rpm. For high density cultivation, strains were grown in Terrific Broth (TB) medium according to reference [80]. For analysis of K5 clinical isolates, strains were cultivated in artificial urine medium according to reference [81]. Cultivation in 96 well plates was carried out in square-well deep well plates shaking at 250 rpm.

The capsule biosynthesis gene cluster (*kps* locus) was deleted from strains using λ red recombination as described in reference [82] with the modification that 5 μM of EDTA was added to the cultures at OD_600_ = 0.3. Primers used are provided in Supplementary Table 2.

For purification or analysis of capsular polysaccharides, biomass was harvested from liquid culture by centrifugation, washed once with water, and cell pellets were stored at −20°C and/or lyophilized prior to processing.

### Release of capsular polysaccharide from cells

Cell pellets were resuspended to an OD_600_ of 150-300 in a final concentration of 50 mM trifluoracetic acid. Hydrolytic release from the cell surface can be conducted at temperatures from 4°C – 80°C over a period from 0.5 – 24 hours. Specific hydrolysis conditions used for each analysis in this study are provided in each figure. Appropriate hydrolysis conditions for each *E. coli* capsule type can be ascertained from Supplementary Data Set 1. Following hydrolysis, samples were cooled to 4°C and clarified by centrifugation at 10’000 g for 15 minutes. The capsule-containing supernatant was then transferred to a new vessel for further purification or analysis.

### Purification of capsular polysaccharides

Capsular polysaccharides were purified from whole-cell hydrolysates as follows. Proteins and lipids were removed by extraction with 5% sample volume of phenol:chloroform:isoamyl alcohol (25:24:1). The aqueous phase was extensively dialysed against 20 mM Tris-HCl pH 7.5 using SpectraPor 12-14 kDa MWCO dialysis tubing. The sample was then adjusted to 200 mM NaCl and 1 mM MgCl_2_, and nucleic acids were digested with DNase I and RNase A.

Capsular polysaccharide was further purified over DEAE Sepharose (∼30 ml column volume) run at 3 ml/min and eluted in a gradient over 15 column volumes from 0 – 1 M NaCl in 20 mM Tris-HCl pH 7.5. Capsule-containing fractions were identified by DMB labelling, dialysed against water, then lyophilized, and resuspended in water at >30 mg/ml. To further reduce contaminants, capsular polysaccharides were precipitated by addition of 5% CTAB, then resuspended in 3 M CaCl_2_, reprecipitated with 3 volumes of ethanol, and finally solubilized in water.

### DMB labelling of capsular polysaccharide samples

Samples containing capsular polysaccharide hydrolysed from the lipid anchor were labelled in DMB labelling reagent composed of 10 mM 1,2-diamino-4,5-methylenedioxybenzene (DMB), 40 mM trifluoracetic acid (TFA), 20 mM sodium dithionite, and 0.5 M β-mercaptoethanol. Labelling reactions proceeded at 25°C for 16-24 hours.

The K1 polysaccharide is subject to formation of reversible intramolecular lactones in acidic conditions which can protect the polysaccharide from excessive hydrolytic degradation [83], [84]. Prior to high resolution anion exchange chromatography, the lactones need to be removed to restore negative charges. To remove lactones, K1 samples were adjusted to 200 mM NaOH and incubated at room temperature for 30 minutes prior to HPLC analysis.

### One-step hydrolysis and DMB labelling

Cell pellets were resuspended to an OD_600_ of 40-80 (2×10^10^ – 4×10^10^ cells/ml) in 100 μl of DMB labelling reagent composed of 10 mM DMB, 40 mM TFA, 20 mM sodium dithionite, and 0.5 M β-mercaptoethanol. From each resuspended pellet, four 20 μl aliquots were transferred to 96-well PCR plates, and each plate incubated for 1 hour at 20°C, 40°C, 60°C, or 80°C, then kept in the dark overnight at 25°C. Each 20 μl labelling reaction was stopped by addition of 100 μl of 20 mM Tris-HCl pH 7.5, then stored at −20°C until HPLC analysis. Prior to injection on the HPLC, samples were clarified by centrifugation at 4000 g for 30 minutes and supernatants transferred to a clean 96 well plate for sampling on the HPLC. 80 μl of each sample was separated by anion exchange chromatography.

Intramolecular lactones were removed from K1 samples prior to HPLC by adjusting to 200 mM NaOH (+ 5 μl of 5 M NaOH) and incubation at room temperature for 30 minutes.

### Anion exchange analysis of capsular polysaccharides

Polysaccharide samples were separated on a DNAPac PA200 column (4 μm, 4.6×150 mm) at 35°C with online fluorescence detection (excitation 373 nm / emission 448 nm). The gradient from 0 – 1 M NaCl (Fig. 1A) was formed by solvent A (20 mM Tris-HCl pH 7.5) and solvent B (1 M NaCl, 20 mM Tris-HCl pH 7.5) as follows: 100% solvent A from 0-2 minutes, curved gradient (Chromeleon gradient No. 3) from 0-40% solvent B from 2-10 minutes, linear gradient from 40-60% solvent B from 10-18 minutes, column wash in 100% solvent B from 18-20 minutes, and re-equilibration in 100% solvent A from 20-27 minutes.

### Bacteriophage-based K5 serotyping assay

K5 serotyping assays were conducted with K5 bacteriophage (Art. No. 60759, SSI Diagnostica). We used the cross-brush method [85] according to the manufacturer’s instructions. A vertical line of phage suspension is applied onto an agar plate by streaking a 10 μl inoculation loop up and down the plate, then left to dry for 10 minutes. A horizontal line of live bacterial culture is streaked across the phage suspension using a 1 μl inoculation loop. The result is evaluated after incubation at 37°C. A positive reaction is indicated by a lack of growth or limited growth after crossing the phage line.

Undisturbed growth along the length of the bacterial streak indicates a negative reaction. Assays were conducted in duplicate for each strain and repeated on two separate occasions.

We also tested phage sensitivity in liquid culture format. LB media was inoculated with 1:100 dilution of dense overnight culture and 150 μl was transferred to the wells of a 96-well plate. Plates were shaken at 37°C in a microtiter plate reader and OD_600_ was measured at 10 minute intervals. At mid-log phase (after ∼150 minutes) the cultures were infected with PBS control, or 10 μl of a 1:100 dilution of K5 bacteriophage stock solution. Assays were conducted in duplicate, using cultures from two separate colonies for each strain.

### K5 lyase reactions

DMB labelled polysaccharides were incubated with purified K5 lyase using established reaction conditions [42]. Reactions proceeded overnight at 37°C in 25 mM Tris-HCl pH 7.5, 50 mM NaCl, 1 mM DTT, with 5 μM of K5 lyase, and no enzyme for control samples. Reactions were diluted with 5 volumes of 20 mM Tris pH 7.5, cleared by centrifugation, and analyzed directly by anion exchange chromatography.

### Genome Sequencing and Bioinformatics Workflow

DNA was recovered from cell pellets by phenol-chloroform extraction. Concentration was measured with a Qubit 2.0 fluorometer and an Agilent Bioanalyzer. Libraries were prepared with the Nextera XT kit and sequenced using paired-end 150 bp reads on the NovaSeq 6000 platform. Raw reads underwent quality control to remove adapters, PhiX contamination, and low-quality bases using bbmap v38.87 [86] with a trimming quality threshold of 14 and a minimum read length of 45 bp.

High-quality reads were assembled using SPAdes v3.14.1 [87]. Capsule gene clusters were identified with fastKaptive [25], and compared using clinker [88].

## Acknowledgments

We thank Andreas Sichert for comments on the manuscript. E.S. and T.G.K. acknowledge a Bridge Discovery Grant (Nr. 180953). T.G.K. is grateful for an SNF Spark Grant (Nr. CRSK-1_190811). Work in this study was funded by the Deutsche Forschungsgemeinschaft (DFG, German Research Foundation)—412824531 to T.F. Work by N.W. and A.B. was supported by SNSF professorships PP00P3_176954 and PP00P3_213978 to M.D. and by the Microbial grant from the Gebert Rüf Stiftung GRS–093/20 to M.D. We acknowledge members of the Swiss Pediatric Sepsis Study for providing the K5 clinical isolates: Philipp Agyeman, Luregn J Schlapbach, Eric Giannoni, Martin Stocker, Klara M Posfay-Barbe, Ulrich Heininger, Sara Bernhard-Stirnemann, Anita Niederer-Loher, Christian Kahlert, Giancarlo Natalucci, Christa Relly, Thomas Riedel, Christoph Aebi, Christoph Berger.

## Conflict of interest

The authors declare no conflict of interest.

## Author contributions

C.R. carried out DMB labelling analyses, purified capsular polysaccharides, constructed capsule deletion mutants, and analyzed data. K.V. and I.D. carried out DMB labelling analyses and optimized hydrolysis conditions. C.M. cloned the K23 capsule gene cluster and conducted experiments to test polysaccharide expression. L.P. optimized DMB labelling reaction conditions. H.L. and C.T. provided unique reagents. M.S. and T.F. provided recombinant K5 lyase. P.S. provided insights into DMB labelling chemistry. L.L., N.W., E.C.B., I.D., A.D., and J.R.N-G. constructed capsule deletion mutants. C.R.M. and S.S. sequenced the genome of clinical isolates and the K5 reference strain and identified kps loci. R.S. carried out K5 lyase reactions and contributed to sequencing of clinical isolates. S.dV. carried out DMB labelling analyses and K5 phage sensitivity assays. Y.T and M.W. contributed to study design. E.S. guided study design. T.G.K. designed the study, conducted some experiments, and wrote the manuscript. All authors read and approved the manuscript.

## Notes

### Competing Interest Statement

The authors have declared no competing interest.

