## Supplementary Tables and Figures for "DMB labelling for detection and analysis of capsular polysaccharides"

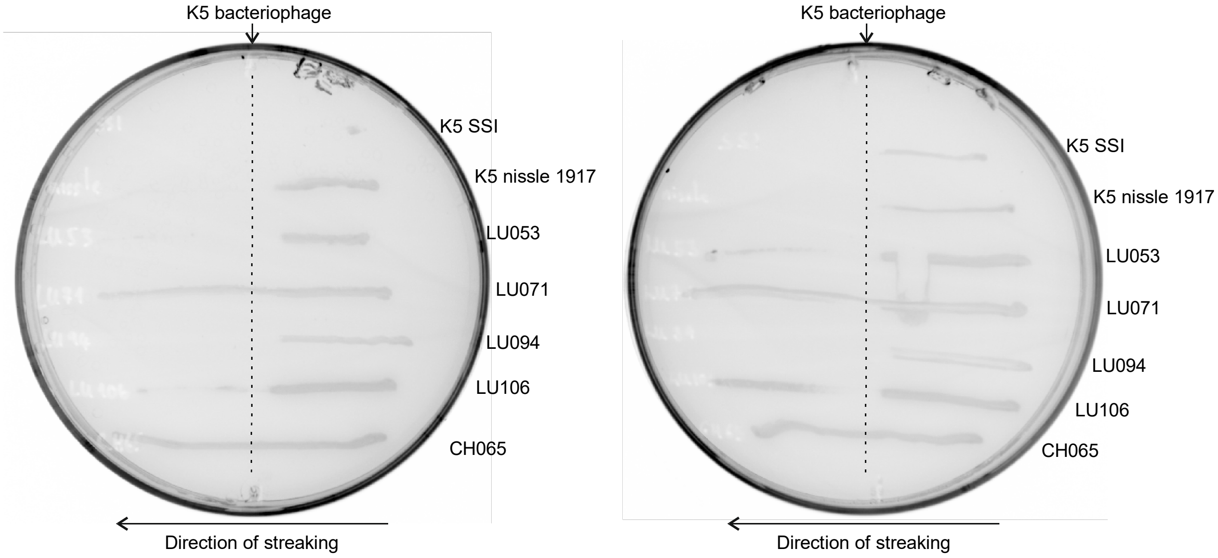


**Supplementary Figure 1. Cross-brush serotyping with K5 bacteriophage.** A vertical line of K5 bacteriophage suspension was applied to the agar plate using an inoculation loop. Live culture from test strains was streaked across the bacteriophage line in the direction indicated. Bacteriophage susceptibility was evaluated after incubation at 37°C. Two replicates are shown.

**Supplementary Table 1. Bacterial strains used in this study**

| Strain name | Source/Reference | K-type | Serotype | Used in |
|---|---|---|---|---|
| E. coli reference strain K1 | SSI 85361 | K1 | O2:K1:H4 | Fig. 3, Extended data Fig. 1 |
| E. coli K2a | CCUG 27 | K2a | O6:K2a:H1 | Fig. 1c & d, Extended data Fig. 1 |
| E. coli reference strain K2ab | SSI 85362 | K2ab | O3:K2ab:H2 | Fig. 1e, Extended data Fig. 1 |
| E. coli K2ab Δkps | derived from SSI 85362 | - | O3:H2 | Fig. 1e |
| E. coli reference strain K3 | SSI 85363 | K3 | O4:K3:H5 | Extended data Fig. 1 |
| E. coli reference strain K4 | SSI 85364 | K4 | O5:K4:H4 | Fig. 1c & d, Fig 3, Extended data Fig. 1 |
| E. coli reference strain K5 | SSI 85365 | K5 | O10:K5:H4 | Fig. 1c & d, Fig. 2, Fig. 3, Extended data Fig. 1 |
| E. coli reference strain K6 | SSI 81964 | K6 | O4:K6:H5 | Fig. 3, Extended data Fig. 1 |
| E. coli reference strain K7 | SSI 81965 | K7 | O7:K7:H4 | Fig. 3, Extended data Fig. 1 |
| E. coli reference strain K8 | SSI 85366 | K8 | O8:K8:H4 | Extended data Fig. 1 |
| E. coli reference strain K9 | SSI 85367 | K9 | O9:K9:H12 | Extended data Fig. 1 |
| E. coli reference strain K10 | SSI 85368 | K10 | O11:K10:H10 | Fig. 3, Extended data Fig. 1 |
| E. coli reference strain K11 | SSI 85369 | K11 | O13:K11:H11 | Fig. 3, Extended data Fig. 1 |
| E. coli reference strain K12 | SSI 81966 | K12 | O4:K12:H- | Fig. 1c & d, Fig. 3, Extended data Fig. 1 |
| E. coli reference strain K13 | SSI 81967 | K13 | O6:K13:H1 | Fig. 1c, d & e, Extended data Fig. 1 |
| E. coli K13 Δkps | derived from SSI 81967 | - | O6:H1 | Fig. 1e |
| E. coli reference strain K14 | SSI 85370 | K14 | O15:K14:H4 | Fig. 1c, d & e, Extended data Fig. 1 |
| E. coli K14 Δkps | derived from SSI 85370 | - | O15:H4 | Fig. 1e |
| E. coli reference strain K16 | SSI 85371 | K16 | O17:K16:H18 | Fig. 3, Extended data Fig. 1 |
| E. coli reference strain K17 | SSI 85372 | K17 | O20:K17:H- | Extended data Fig. 1 |
| E. coli reference strain K18a | SSI 81968 | K18a | O23:K18a:H15 | Extended data Fig. 1 |
| E. coli reference strain K18ab | SSI 85373 | K18ab | O23:K18ab:H15 | Extended data Fig. 1 |
| E. coli reference strain K19 | SSI 85374 | K19 | O25:K19:H12 | Fig. 3, Extended data Fig. 1 |
| E. coli reference strain K20 | SSI 85375 | K20 | O21:K20:H- | Fig. 3, Extended data Fig. 1 |
| E. coli K22 | CCUG 11329 | K22 | O23:K22:H15 | Extended data Fig. 1 |
| E. coli reference strain K23 | SSI 81970 | K23 | O25:K23:H1 | Fig. 3, Extended data Fig. 1 |
| E. coli K23 | NCTC 10430 | K23 | O25:K23:H1 | Fig. 1e |
| E. coli K23 Δkps | derived from NCTC 10430 | - | O25:H1 | Fig. 1e |
| E. coli reference strain K24 | SSI 81971 | K24 | O83:K24:H31 | Fig. 3, Extended data Fig. 1 |
| E. coli reference strain K26 | SSI 81972 | K26 | O9:K26:H- | Extended data Fig. 1 |
| E. coli reference strain K27 | SSI 81973 | K27 | O8:K27:H- | Extended data Fig. 1 |
| E. coli reference strain K28 | SSI 81974 | K28 | O9:K28:H- | Extended data Fig. 1 |
| E. coli reference strain K29 | SSI 81975 | K29 | O9:K29:H- | Extended data Fig. 1 |
| E. coli reference strain K30 | SSI 81976 | K30 | O9:K30:H12 | Extended data Fig. 1 |
| E. coli reference strain K31 | SSI 81977 | K31 | O9:K31:H- | Extended data Fig. 1 |
| E. coli reference strain K32 | SSI 81978 | K32 | O9:K32:H19 | Extended data Fig. 1 |
| E. coli reference strain K33 | SSI 81979 | K33 | O9:K33:H- | Extended data Fig. 1 |
| E. coli reference strain K34 | SSI 81980 | K34 | O9:K34:H- | Extended data Fig. 1 |
| E. coli reference strain K35 | SSI 81981 | K35 | O9:K35:H- | Extended data Fig. 1 |
| E. coli reference strain K36 | SSI 81982 | K36 | O9:K36:H19 | Extended data Fig. 1 |
| E. coli reference strain K37 | SSI 81983 | K37 | O9:K37:H- | Extended data Fig. 1 |
| E. coli reference strain K38 | SSI 81984 | K38 | O9:K38:H- | Extended data Fig. 1 |
| E. coli reference strain K39 | SSI 81985 | K39 | O9:K39:H9 | Extended data Fig. 1 |
| E. coli reference strain K40 | SSI 81986 | K40 | O8:K40:H9 | Extended data Fig. 1 |
| E. coli reference strain K41 | SSI 81987 | K41 | O8:K41:H11 | Extended data Fig. 1 |
| E. coli reference strain K42 | SSI 81988 | K42 | O8:K42:H- | Extended data Fig. 1 |
| E. coli reference strain K43 | SSI 81989 | K43 | 08:K43:H11 | Extended data Fig. 1 |
| E. coli reference strain K44 | SSI 81990 | K44 | O8:K44:H- | Extended data Fig. 1 |
| E. coli reference strain K45 | SSI 81991 | K45 | O8:K45:H9 | Extended data Fig. 1 |
| E. coli reference strain K46 | SSI 81992 | K46 | O8:K46:K30 | Extended data Fig. 1 |
| E. coli reference strain K47 | SSI 81993 | K47 | O8:K47:H2 | Extended data Fig. 1 |
| E. coli reference strain K48 | SSI 81994 | K48 | O8:K48:H9 | Extended data Fig. 1 |
| E. coli reference strain K49 | SSI 81995 | K49 | O8:K49:H21 | Extended data Fig. 1 |
| E. coli reference strain K50 | SSI 81996 | K50 | O8:K50:H- | Extended data Fig. 1 |
| E. coli reference strain K51 | SSI 81997 | K51 | O1:K51:H- | Extended data Fig. 1 |
| E. coli reference strain K52 | SSI 81998 | K52 | O4:K52:H- | Extended data Fig. 1 |
| E. coli reference strain K53 | SSI 81999 | K53 | O6:K53:H- | Extended data Fig. 1 |
| E. coli reference strain K54 | SSI 82000 | K54 | O6:K54:H10 | Extended data Fig. 1 |
| E. coli reference strain K55 | SSI 82001 | K55 | O9:K55:H- | Extended data Fig. 1 |
| E. coli reference strain K83 | SSI 82003 | K83 | O20:K83:H26 | Extended data Fig. 1 |
| E. coli reference strain K87 | SSI 82005 | K87 | O8:K87:H19 | Extended data Fig. 1 |
| E. coli reference strain K92 | SSI 82006 | K92 | O73:K92:H34 | Extended data Fig. 1 |
| E. coli reference strain K93 | SSI 85376 | K93 | O150:K93:H6 | Extended data Fig. 1 |
| E. coli reference strain K95 | SSI 85378 | K95 | O75:K95:H5 | Extended data Fig. 1 |
| E. coli reference strain K96 | SSI 85379 | K96 | O77:K96:H- | Fig. 3, Extended data Fig. 1 |
| E. coli reference strain K97 | SSI 85380 | K97 | O81:K97:H- | Fig. 3, Extended data Fig. 1 |
| E. coli reference strain K99 | SSI 85381 | K98 | O107:K98:H27 | Fig. 3, Extended data Fig. 1 |
| E. coli reference strain K100 | SSI 82007 | K100 | O75:K100:H5 | Extended data Fig. 1 |
| E. coli reference strain K101 | SSI 82008 | K101 | O20:K101:H-:F6 | Extended data Fig. 1 |
| E. coli reference strain K102 | SSI 82009 | K102 | O8:K102:H- | Extended data Fig. 1 |
| E. coli reference strain K103 | SSI 82010 | K103 | O101:K103:H- | Extended data Fig. 1 |
| CH065 | Agyeman *et al.,* (2017) | - | | Fig. 2 |
| LU053 | Agyeman *et al.,* (2017) | K5 | | Fig. 1e, Fig. 2 |
| E. coli K5 Δkps | derived from LU053 | - | | Fig. 1e |
| LU106 | Agyeman *et al.,* (2017) | K5 | | Fig. 2 |
| LU094 | Agyeman *et al.,* (2017) | K5 | | Fig. 2 |
| LU071 | Agyeman *et al.,* (2017) | K5 | | Fig. 2 |
| LU100 | Agyeman *et al.,* (2017) | K1 | | Fig. 1e |
| E. coli K1 Δkps | derived from LU100 | - | | Fig. 1e |
| E. coli Nissle 1917 | Institute of Microbiology, ETHZ | K5 | O6:K5:H1 | Fig. 2 |
| E. coli HS | Levine, Sotman et al., Lancet (1978) | | O9:H4 | Extended data Fig. 1 |
| E. coli W3110 | CGSC #4474 | | | Extended data Fig. 1 |

**Supplementary Table 2. Primers**

| Purpose | Sequence (5' - 3') |
|---|---|
| Fwd primer for pKD3/pKD4, used for K1, K5, & K23 Δkps | ATGTCTGAAAGACATTTACCTGATGACCAGAGCAGTACTATGTAGGCTGGAGCTGCTTCG |
| Rev primer for pKD3/pDK4, used for K1, K5, & K23 Δkps | TTGGTAGCTGTTAAGCCAAGGGCGGTAGCGTACCTGAAGAATGGGAATTAGCCATGGTCC |
| Fwd primer for pKD3, deletion: K2ab, K13, & K14 Δkps | ATGTCTGAAAGACATTTACCTGATGACCAGAGCAGTACTATGTAGGCTGGAGCTGCTTC |
| Rev primer for pKD3, used for K2ab Δkps | TTGGTAGCTGTTAAGCCAGGGGCGGTAGCATACCTGAAGACATATGAATATCCTCCTTAGTT |
| Rev primer for pKD3, used for K13 Δkps | TTGGTAGCTGTTAAGCCAAGGGCGGTAGCGTACCTGAAGACATATGAATATCCTCCTTAGTT |
| Rev primer for pKD3, used for K14 Δkps | TTGGTAGCTGTTAAGCCAGGGGCGGTAGCGTACCTGAAGACATATGAATATCCTCCTTAGTT |

**Negative control - E. coli W3110**

**Capsule type:** none
**Strain**: *E. coli* K-12 wild-type W3110
**Strain No.**: CGSC #4474
**Reference**: Jensen, J Bacteriol, (1993)
**Culture media**: TB
**Approx. per injection**: 1.6e9 cells

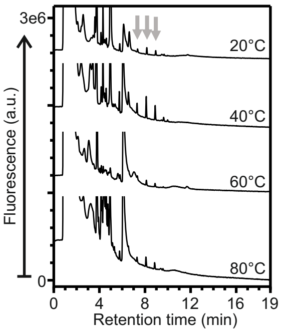


**Negative control - E. coli HS**

**Capsule type:** none
**Strain**: *E. coli* HS
**Reference**: Levine, Sotman, *et al.*, Lancet, (1978)
**Culture media**: LB
**Approx. per injection**: 2e8 cells

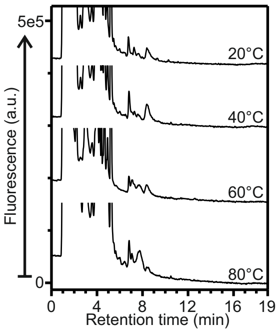


**Supplementary Data Set 1. DMB-labelling of E. coli K antigen reference strains.** A comprehensive collection of *E. coli* strains expressing the full spectrum of capsular polysaccharides was analyzed by DMB labelling and anion exchange chromatography.  Strains were cultivated in 96-well plates in LB and/or TB media. The harvested cell pellets were washed with water, lyophilized, and resuspended in DMB labelling reagent. Hydrolysis of the polysaccharide and DMB labeling was allowed to proceeded in parallel. To account for different hydrolytic stability of the polysaccharides, samples were incubated at 20°C, 40°C, 60°C, or 80°C. Supernatants were analysed by anion exchange chromatography with online fluorescence detection.

Page 1: Unencapsulated strains. Negative controls.
Page 2-4: Transporter-dependent capsular polysaccharides (group 2 & 3 capsules)
Page 5-7: Wzx-Wzy-dependent capsular polysaccharides (group 1 & 4 capsules). Specificity controls.

#### Legend

- putative capsular polysaccharide
- autofluorescent species (observed without DMB-labelling)
- glucose (Glc)
- galactose (Gal)
- 3-deoxy-D-manno-oct-2-ulosonic acid (Kdo)
- N-acetylglucosamine (GlcNAc)
- galactofuranose (Gal*f*)
- 4,6-dideoxy-4-malonylaminoglucose (QuiNMal)
- glucuronic acid (GlcA)
- N-acetylgalactosamine (GalNAc)
- N-acetylmannosaminuronic acid (ManNAcA)
- rhamnose (Rha)
- sialic acid (Neu5Ac)
- 4-deoxy-2-hexulosonic acid
- fructofuranose (Fruc*f*)
- ribofuranose (Rib*f*)

| | |
|---|---|
| Gro | glycerol |
| Rib | ribitol |
| P | phosphate |
| Ac | acetyl |
| Prop | propionyl |

### Supplementary Data Set - Page 2

**K1 repeat unit**:

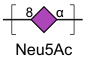


**Capsule type:** group 2
transporter-dependent
**Strain**: *E. coli* reference strain K1
**Strain No.**: SSI #85361
**Serotype**: O2:K1:H4
**Reference**: McGuire and Blinkley, Biochemistry, (1964)
**Culture media**: LB
**Approx. per injection**: 2e8 cells

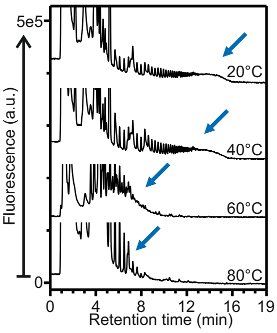


**K2a**:

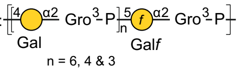


**Capsule type:** group 2
transporter-dependent
**Strain**: *E. coli* Bi 7458-41
**Strain No.**: CCUG #27
**Serotype**: O6:K2a:H1
**Reference**: Fischer, Schmidt, Jann and Jann, Biochemistry, (1982)
**Culture media**: TB
**Approx. per injection**: 1.6e9 cells

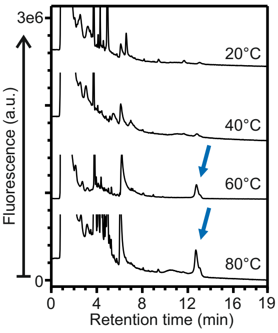


**K2ab**:

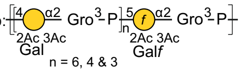


**Capsule type:** group 2
transporter-dependent
**Strain**: *E. coli* reference strain K2ab
**Strain No.**: SSI #85362
**Serotype**: O3:K2ab:H2
**Reference**: Fischer, Schmidt, Jann and Jann, Biochemistry, (1982)
**Culture media**: TB
**Approx. per injection**: 1.6e9 cells

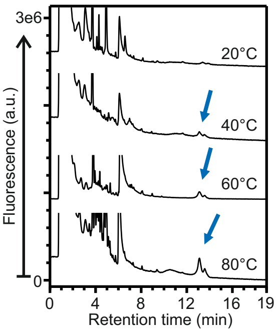


**K3 repeat unit**:

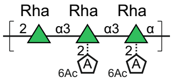


**Capsule type:** group 3
transporter-dependent
**Strain**: *E. coli* reference strain K3
**Strain No.**: SSI #85363
**Serotype**: O4:K3:H5
**Reference**: Dengler, Jann and Jann, Carb. Res., (1988)
**Culture media**: TB
**Approx. per injection**: 1.6e9 cells

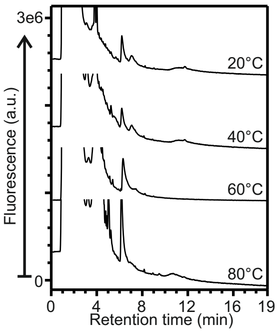


**K4 repeat unit**:

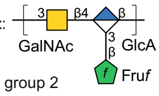


**Capsule type:** group 2
transporter-dependent
**Strain**: *E. coli* reference strain K4
**Strain No.**: SSI #85364
**Serotype**: O5:K4:H4
**Reference**: Rodriguez, Jann and Jann, Eur J Biochem, (1988)
**Culture media**: LB
**Approx. per injection**: 2e8 cells

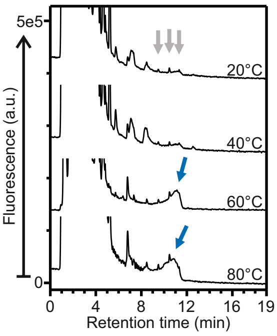


**K5 repeat unit**:

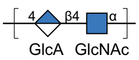


**Capsule type:** group 2
transporter-dependent
**Strain**: *E. coli* reference strain K5
**Strain No.**: SSI #85365
**Serotype**: O10:K5:H4
**Reference**: Vann, Schmidt, Jann and Jann, Eur J Biochem, (1981)
**Culture media**: TB
**Approx. per injection**: 1.6e9 cells

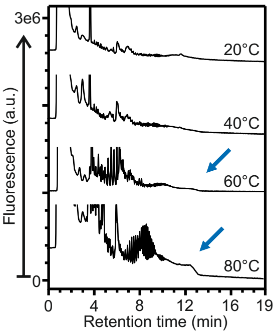


**K6 repeat unit**:

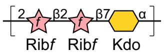


**Capsule type:** group 2
transporter-dependent
**Strain**: *E. coli* reference strain K6
**Strain No.**: SSI #81964
**Serotype**: O4:K6:H5
**Reference**: Jennings, Rosell and Johnson Carb Res (1982)
**Culture media**: LB
**Approx. per injection**: 2e8 cells

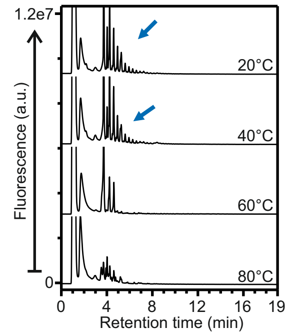


**K7/K56 repeat unit**:

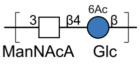


**Capsule type:** group 2
transporter-dependent
**Strain**: *E. coli* reference strain K7
**Strain No.**: SSI #81965
**Serotype**: O7:K7:H4
**Reference**: Tsui, Boykins and Egan, Carb Res (1982)
**Culture media**: LB
**Approx. per injection**: 2e8 cells

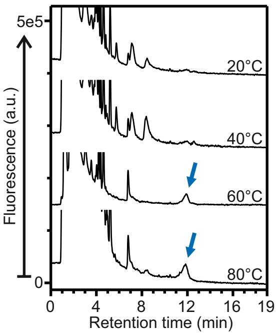


**K10 repeat unit**:

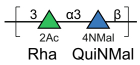


**Capsule type:** group 3
transporter-dependent
**Strain**: *E. coli* reference strain K10
**Strain No.**: SSI #85368
**Serotype**: O11:K10:H10
**Reference**: Sieberth, Jann and Jann, Carb Res, (1993)
**Culture media**: TB
**Approx. per injection**: 1.6e9 cells

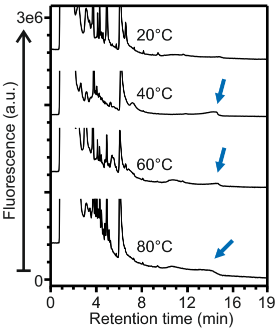


**K11 repeat unit**:

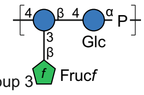


**Capsule type:** group 3
transporter-dependent
**Strain**: *E. coli* reference strain K11
**Strain No.**: SSI #85369
**Serotype**: O13:K11:H11
**Reference**: Rodriguez, Jann and Jann, Carb Res, (1990)
**Culture media**: TB
**Approx. per injection**: 1.6e9 cells

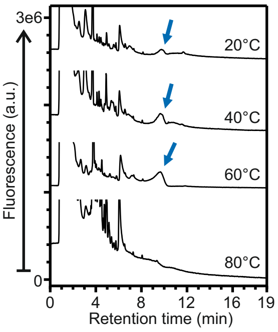


**K12 repeat unit**:

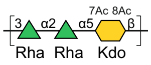


**Capsule type:** group 2
transporter-dependent
**Strain**: *E. coli* reference strain K12
**Strain No.**: SSI #81966
**Serotype**: O4:K12:H-
**Reference**: Schmidt and Jann, Eur J Biochem, (1983)
**Culture media**: TB
**Approx. per injection**: 1.6e9 cells

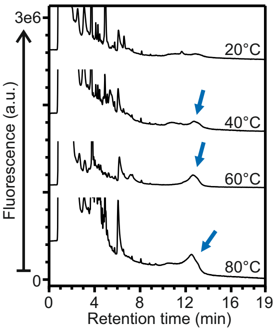


**K13 repeat unit**:

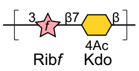


**Capsule type:** group 2
transporter-dependent
**Strain**: *E. coli* reference strain K13
**Strain No.**: SSI #81967
**Serotype**: O6:K13:H1
**Reference**: Vann and Jann, Infect Immun, (1979)
**Culture media**: TB
**Approx. per injection**: 1.6e9 cells

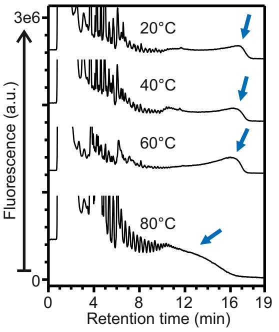

**K14 repeat unit**:

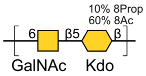


**Capsule type:** group 2
transporter-dependent
**Strain**: *E. coli* reference strain K14
**Strain No.**: SSI #85370
**Serotype**: O15:K14:H4
**Reference**: Jann, Hofman and Jann
Carb Res, (1983)
**Culture media**: TB
**Approx. per injection**: 1.6e9 cells

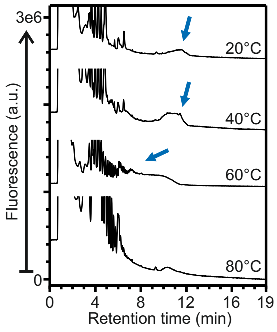


**K16 repeat unit**:

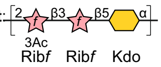


**Capsule type:** group 2
transporter-dependent
**Strain**: *E. coli* reference strain K16
**Strain No.**: SSI #85371
**Serotype**: O17:K16:H18
**Reference**: Lenter, Jann and Jann,
Carb Res, (1990)
**Culture media**: TB
**Approx. per injection**: 1.6e9 cells

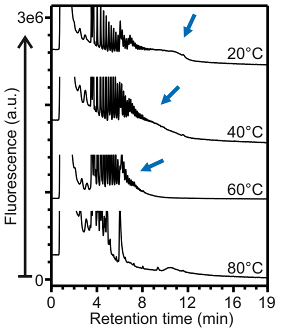


**K18a repeat unit**:

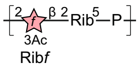


**Capsule type:** group 2
transporter-dependent
**Strain**: *E. coli* reference strain K18a
**Strain No.**: SSI #81968
**Serotype**: O23:K18a:H15
**Reference**: Rodriguez, Jann and Jann,
Carb Res, (1988)
**Culture media**: TB
**Approx. per injection**: 1.6e9 cells

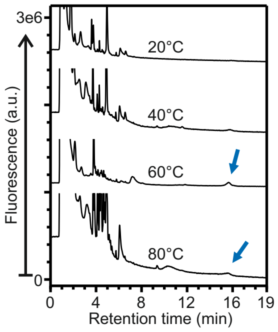


**K18ab repeat unit**:

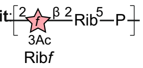


**Capsule type:** group 2
transporter-dependent
**Strain**: *E. coli* reference strain K18ab
**Strain No.**: SSI #85373
**Serotype**: O23:K18ab:H15
**Reference**: Rodriguez, Jann and Jann,
Carb Res, (1988)
**Culture media**: TB
**Approx. per injection**: 1.6e9 cells

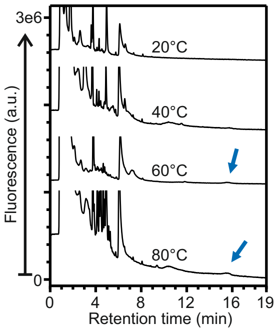


**K19 repeat unit**:

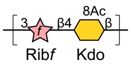


**Capsule type:** group 2
transporter-dependent
**Strain**: *E. coli* reference strain K19
**Strain No.**: SSI #85374
**Serotype**: O25:K19:H12
**Reference**: Jann, Ahrens, Dengler and
Jann, Carb Res, (1988)
**Culture media**: LB
**Approx. per injection**: 2e8 cells

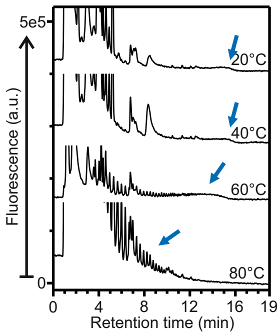


**K20 repeat unit**:

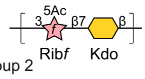


**Capsule type:** group 2
transporter-dependent
**Strain**: *E. coli* reference strain K20
**Strain No.**: SSI #85375
**Serotype**: O21:K20:H-
**Reference**: Vann, Soderstrom, Egan,
Tsui, Schneerson, Ørskov and Ørskov,
Infect Immun, (1983)
**Culture media**: TB
**Approx. per injection**: 1.6e9 cells

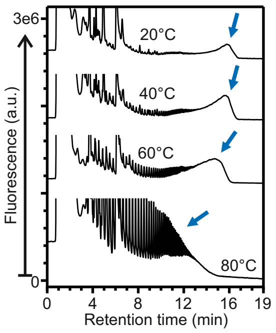


**K22 repeat unit**:

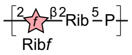


**Capsule type:** group 2
transporter-dependent
**Strain**: *E. coli* K22
**Strain No.**: CCUG #11329
**Serotype**: O23:K22:H15
**Reference**: Rodriguez, Jann and Jann
Carb Res, (1988)
**Culture media**: TB
**Approx. per injection**: 1.6e9 cells

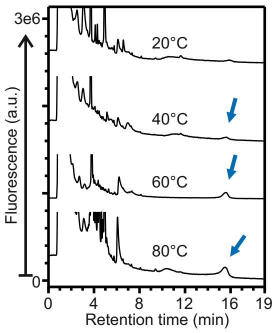


**K23 repeat unit**:

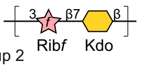


**Capsule type:** group 2
transporter-dependent
**Strain**: *E. coli* reference strain K23
**Strain No.**: SSI #81970
**Serotype**: O25:K23:H1
**Reference**: Vann, Soderstrom, Egan,
Tsui, Schneerson, Ørskov and Ørskov,
Infect Immun, (1983)
**Culture media**: LB
**Approx. per injection**: 2e8 cells

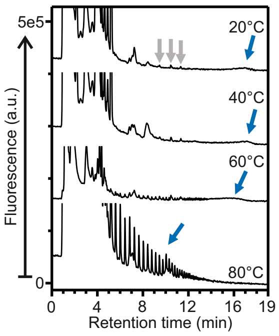


**K24 repeat unit**:

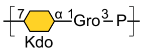


**Capsule type:** group 2
transporter-dependent
**Strain**: *E. coli* reference strain K24
**Strain No.**: SSI #81971
**Serotype**: O83:K24:H31
**Reference**: Lenter, Jann and Jann
Carb Res, (1990)
**Culture media**: TB
**Approx. per injection**: 1.6e9 cells

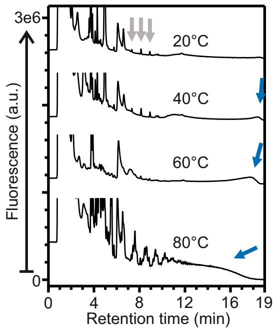


**K51 repeat unit**:

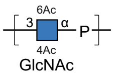


**Capsule type:** group 2
transporter-dependent
**Strain**: *E. coli* reference strain K51
**Strain No.**: SSI #81997
**Serotype**: O1:K51:H-
**Reference**: Jann, Dengler and Jann
FEMS Micro Lett, (1985)
**Culture media**: TB
**Approx. per injection**: 1.6e9 cells

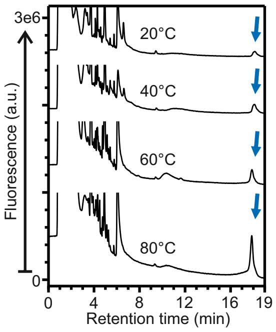


**K52 repeat unit**:

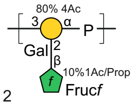


**Capsule type:** group 2
transporter-dependent
**Strain**: *E. coli* reference strain K52
**Strain No.**: SSI #81998
**Serotype**: O4:K52:H-
**Reference**: Hofmann, Jann and Jann
Eur J Biochem, (1985)
**Culture media**: TB
**Approx. per injection**: 1.6e9 cells

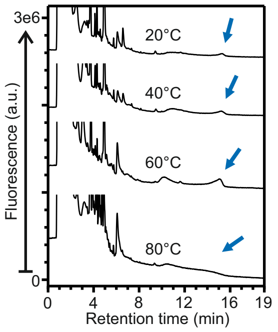


**K53 repeat unit**:

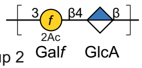


**Capsule type:** group 2
transporter-dependent
**Strain**: *E. coli* reference strain K53
**Strain No.**: SSI #81999
**Serotype**: O6:K53:H-
**Reference**: Bax, Summers, Egan, Guiris
Schneerson, Ørskov, Ørskov and Vann
Carb Res, (1988)
**Culture media**: TB
**Approx. per injection**: 1.6e9 cells

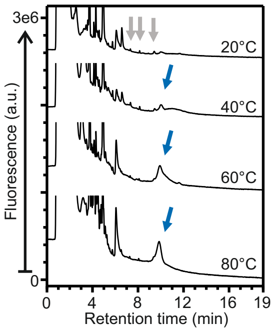

**K54 repeat unit**: 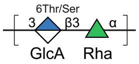


**Capsule type:** group 3
transporter-dependent
**Strain**: *E. coli* reference strain K54
**Strain No.**: SSI #82000
**Serotype**: O6:K54:H10
**Reference**: Jann, Kochanowski and Jann, Carb Res (1994)
**Culture media**: TB
**Approx. per injection**: 1.6e9 cells

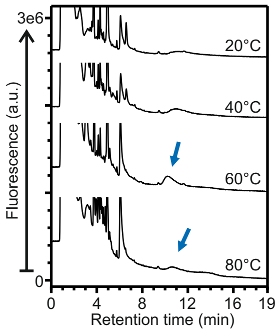


**K92 repeat unit**: 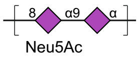


**Capsule type:** group 2
transporter-dependent
**Strain**: *E. coli* reference strain K92
**Strain No.**: SSI #82006
**Serotype**: O73:K92:H34
**Reference**: Egan, Gotschlich and Robbins, *et al.*, Biochemistry, (1977)
**Culture media**: LB
**Approx. per injection**: 2e8 cells

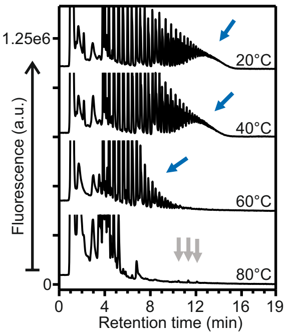


**K93 repeat unit**: 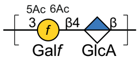


**Capsule type:** group 2
transporter-dependent
**Strain**: *E. coli* reference strain K93
**Strain No.**: SSI #85376
**Serotype**: O150:K93:H6
**Reference**: Bax, Summers, Egan, Guirgis, Schneerson, Robbins, Orskov, Orskov, and Vann, Carb Res (1988)
**Culture media**: TB
**Approx. per injection**: 1.6e9 cells

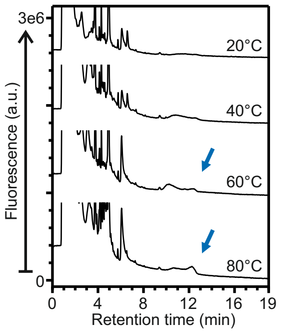


**K95 repeat unit**: 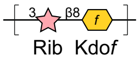


**Capsule type:** group 2
transporter-dependent
**Strain**: *E. coli* reference strain K95
**Strain No.**: SSI #85378
**Serotype**: O75:K95:H5
**Reference**: Dengler, Jann and Jann Biochemistry, (1977)
**Culture media**: LB
**Approx. per injection**: 2e8 cells

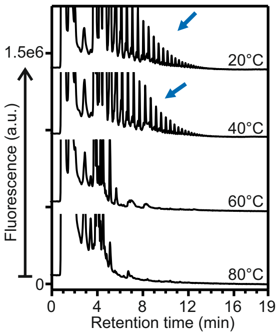


**K96 repeat unit**: 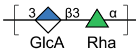


**Capsule type:** group 3
transporter-dependent
**Strain**: *E. coli* reference strain K96
**Strain No.**: SSI #85379
**Serotype**: O77:K96:H-
**Reference**: Hofmann, Jann and Jann Biochemistry, (1985)
**Culture media**: LB
**Approx. per injection**: 2e8 cells

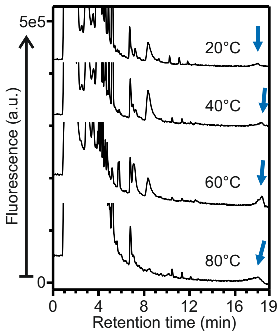


**K97 repeat unit**: 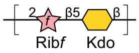


**Capsule type:** group 2
transporter-dependent
**Strain**: *E. coli* reference strain K97
**Strain No.**: SSI #85380
**Serotype**: O81:K97:H-
**Reference**: Jann and Jann,
Curr Top Microbiol Immunol, (1990)
**Culture media**: LB
**Approx. per injection**: 2e8 cells

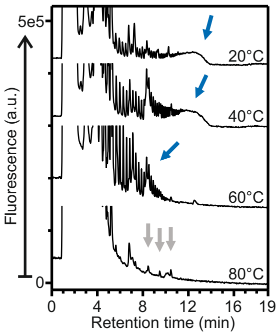


**K98 repeat unit**: 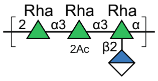


**Capsule type:** group 3
transporter-dependent
**Strain**: *E. coli* reference strain K98
**Strain No.**: SSI #85381
**Serotype**: O107:K98:H27
**Reference**: Hanne, Jann and Jann Carb Res, (1991)
**Culture media**: TB
**Approx. per injection**: 1.6e9 cells

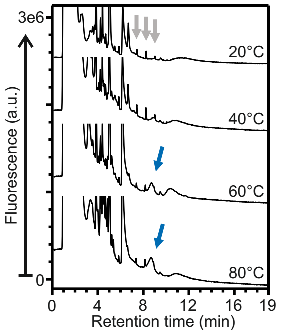


**K100 repeat unit**: 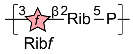


**Capsule type:** group 2
transporter-dependent
**Strain**: *E. coli* reference strain 100
**Strain No.**: SSI #82007
**Serotype**: O75:K100:H5
**Reference**: Rodriguez, Jann and Jann Carb Res (1988)
**Culture media**: TB
**Approx. per injection**: 1.6e9 cells

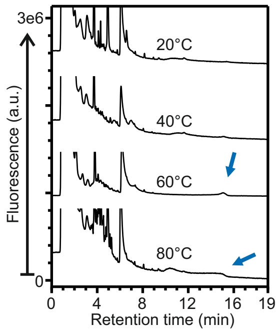

**Negative control - K8**

**Capsule type:** group 4
Wzx/Wzy-dependent
**Strain**: *E. coli* reference strain K8
**Strain No.**: SSI #85366
**Serotype**: O8:K8:H4
**Culture media**: TB
**Approx. per injection**: 1.6e9 cells

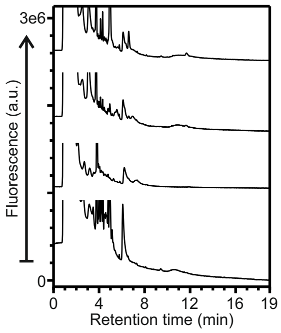


**Negative control - K9**

**Capsule type:** group 4
Wzx/Wzy-dependent
**Strain**: *E. coli* reference strain K9
**Strain No.**: SSI #85367
**Serotype**: O9:K9:H12
**Culture media**: LB
**Approx. per injection**: 2e8 cells

**K9 repeat unit:**

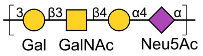


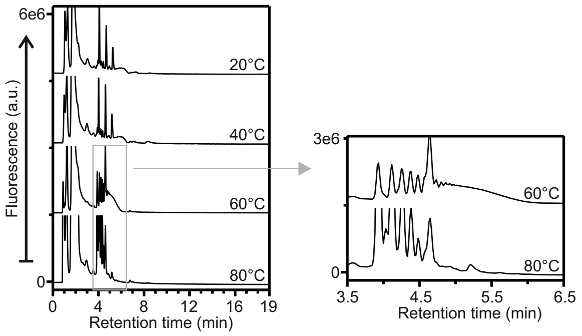


**Negative control - K26**

**Capsule type:** group 1
Wzx/Wzy-dependent
**Strain**: *E. coli* reference strain K26
**Strain No.**: SSI #81972
**Serotype**: O9:K26:H-
**Culture media**: TB
**Approx. per injection**: 1.6e9 cells

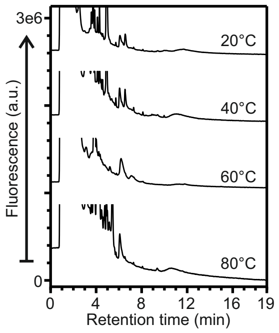


**Negative control - K27**

**Capsule type:** group 1
Wzx/Wzy-dependent
**Strain**: *E. coli* reference strain K27
**Strain No.**: SSI #81973
**Serotype**: O8:K27:H-
**Culture media**: TB
**Approx. per injection**: 1.6e9 cells

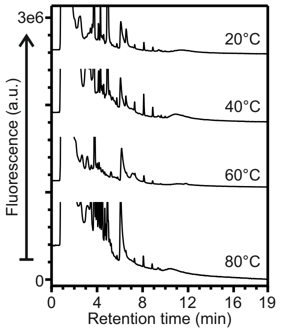


**Negative control - K28**

**Capsule type:** group 1
Wzx/Wzy-dependent
**Strain**: *E. coli* reference strain K28
**Strain No.**: SSI #81974
**Serotype**: O9:K28:H-
**Culture media**: TB
**Approx. per injection**: 1.6e9 cells

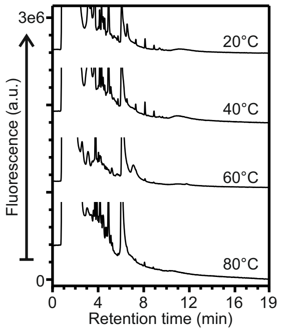


**Negative control - K29**

**Capsule type:** group 1
Wzx/Wzy-dependent
**Strain**: *E. coli* reference strain K29
**Strain No.**: SSI #81975
**Serotype**: O9:K29:H-
**Culture media**: TB
**Approx. per injection**: 1.6e9 cells

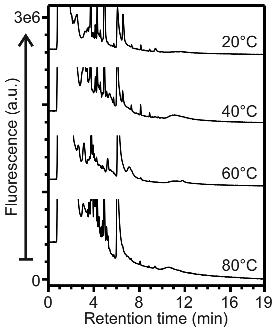


**Negative control - K30**

**Capsule type:** group 1
Wzx/Wzy-dependent
**Strain**: *E. coli* reference strain K30
**Strain No.**: SSI #81976
**Serotype**: O9:K30:H12
**Culture media**: TB
**Approx. per injection**: 1.6e9 cells

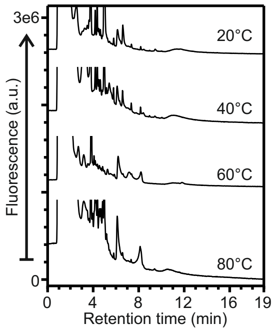


**Negative control - K31**

**Capsule type:** group 1
Wzx/Wzy-dependent
**Strain**: *E. coli* reference strain K31
**Strain No.**: SSI #81977
**Serotype**: O9:K31:H-
**Culture media**: TB
**Approx. per injection**: 1.6e9 cells

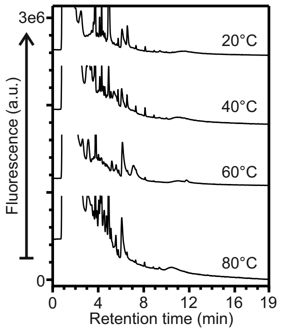


**Negative control - K32**

**Capsule type:** group 1
Wzx/Wzy-dependent
**Strain**: *E. coli* reference strain K32
**Strain No.**: SSI #81978
**Serotype**: O9:K32:H19
**Culture media**: LB
**Approx. per injection**: 2e8 cells

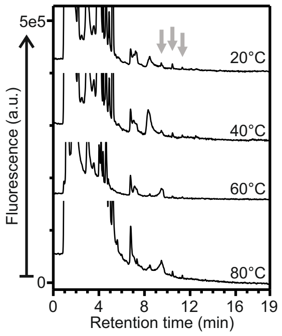


**Negative control - K33**

**Capsule type:** group 1
Wzx/Wzy-dependent
**Strain**: *E. coli* reference strain K33
**Strain No.**: SSI #81979
**Serotype**: O9:K33:H-
**Culture media**: LB
**Approx. per injection**: 2e8 cells

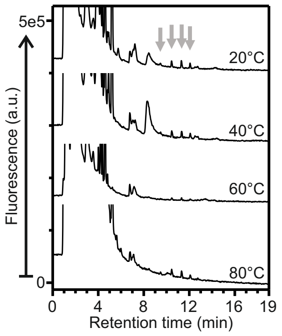


**Negative control - K34**

**Capsule type:** group 1
Wzx/Wzy-dependent
**Strain**: *E. coli* reference strain K34
**Strain No.**: SSI #81980
**Serotype**: O9:K34:H-
**Culture media**: LB
**Approx. per injection**: 2e8 cells

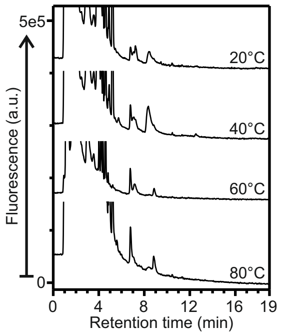

**Negative control - K35**

**Capsule type:** group 1
Wzx/Wzy-dependent
**Strain**: *E. coli* reference strain K35
**Strain No.**: SSI #81981
**Serotype**: O9:K35:H-
**Culture media**: LB
**Approx. per injection**: 2e8 cells

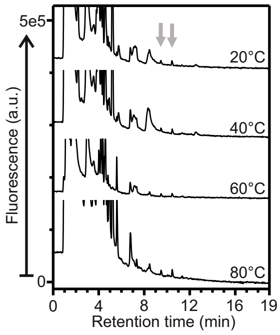


**Negative control - K36**

**Capsule type:** group 1
Wzx/Wzy-dependent
**Strain**: *E. coli* reference strain K36
**Strain No.**: SSI #81982
**Serotype**: O9:K36:H19
**Culture media**: LB
**Approx. per injection**: 2e8 cells

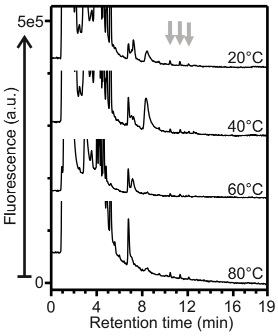


**Negative control - K37**

**Capsule type:** group 1
Wzx/Wzy-dependent
**Strain**: *E. coli* reference strain K37
**Strain No.**: SSI #81983
**Serotype**: O9:K37:H-
**Culture media**: LB
**Approx. per injection**: 2e8 cells

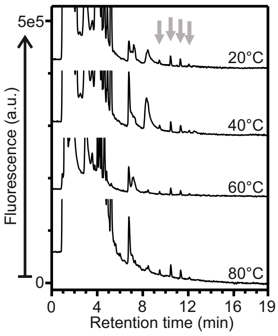


**Negative control - K38**

**Capsule type:** group 4
Wzx/Wzy-dependent
**Strain**: *E. coli* reference strain K38
**Strain No.**: SSI #81984
**Serotype**: O9:K38:H-
**Culture media**: LB
**Approx. per injection**: 2e8 cells

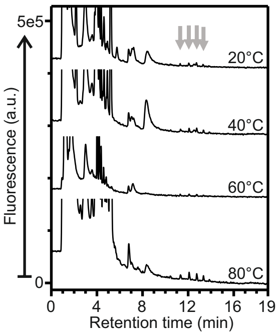


**Negative control - K39**

**Capsule type:** group 1
Wzx/Wzy-dependent
**Strain**: *E. coli* reference strain K39
**Strain No.**: SSI #81985
**Serotype**: O9:K39:H9
**Culture media**: LB
**Approx. per injection**: 2e8 cells

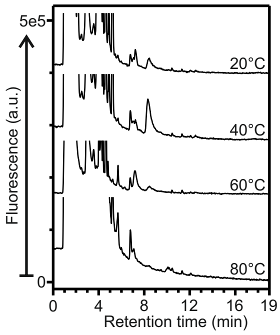


**Negative control - K40**

**Capsule type:** group 4
Wzx/Wzy-dependent
**Strain**: *E. coli* reference strain K40
**Strain No.**: SSI #81986
**Serotype**: O8:K40:H9
**Culture media**: LB
**Approx. per injection**: 2e8 cells

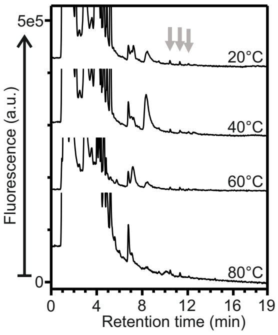


**Negative control - K41**

**Capsule type:** group 1
Wzx/Wzy-dependent
**Strain**: *E. coli* reference strain K41
**Strain No.**: SSI #81987
**Serotype**: O8:K41:H11
**Culture media**: LB
**Approx. per injection**: 2e8 cells

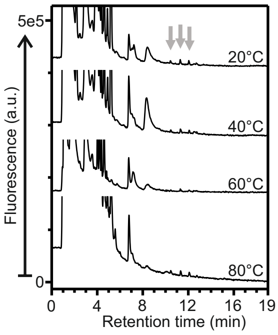


**Negative control - K42**

**Capsule type:** group 1
Wzx/Wzy-dependent
**Strain**: *E. coli* reference strain K42
**Strain No.**: SSI #81988
**Serotype**: O8:K42:H-
**Culture media**: LB
**Approx. per injection**: 2e8 cells

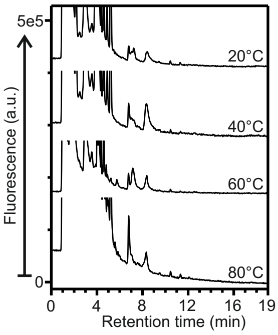


**Negative control - K43**

**Capsule type:** group 1
Wzx/Wzy-dependent
**Strain**: *E. coli* reference strain K43
**Strain No.**: SSI #81989
**Serotype**: 08:K43:H11
**Culture media**: LB
**Approx. per injection**: 2e8 cells

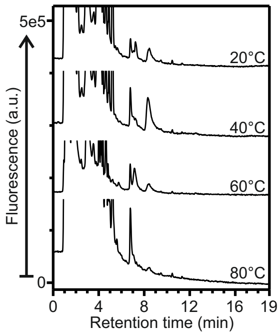


**Negative control - K44**

**Capsule type:** group 4
Wzx/Wzy-dependent
**Strain**: *E. coli* reference strain K44
**Strain No.**: SSI #81990
**Serotype**: O8:K44:H-
**Culture media**: LB
**Approx. per injection**: 2e8 cells

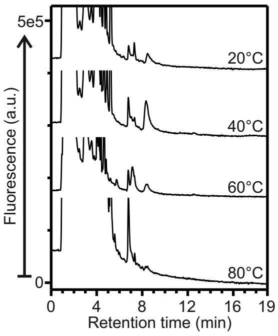


**Negative control - K45**

**Capsule type:** group 4
Wzx/Wzy-dependent
**Strain**: *E. coli* reference strain K45
**Strain No.**: SSI #81991
**Serotype**: O8:K45:H9
**Culture media**: LB
**Approx. per injection**: 2e8 cells

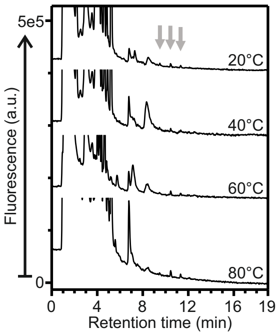


**Negative control - K46**

**Capsule type:** group 4
Wzx/Wzy-dependent
**Strain**: *E. coli* reference strain K46
**Strain No.**: SSI #81992
**Serotype**: O8:K46:K30
**Culture media**: LB
**Approx. per injection**: 2e8 cells

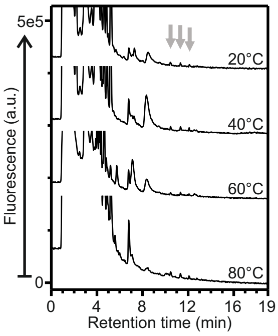

**Negative control - K47**

**Capsule type:** group 4
Wzx/Wzy-dependent
**Strain**: *E. coli* reference strain K47
**Strain No.**: SSI #81993
**Serotype**: O8:K47:H2
**Culture media**: LB
**Approx. per injection**: 2e8 cells

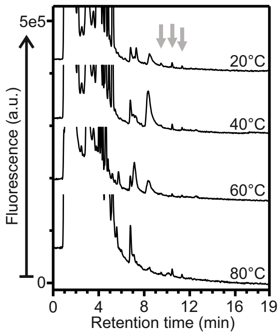


**Negative control - K48**

**Capsule type:** group 4
Wzx/Wzy-dependent
**Strain**: *E. coli* reference strain K48
**Strain No.**: SSI #81994
**Serotype**: O8:K48:H9
**Culture media**: LB
**Approx. per injection**: 2e8 cells

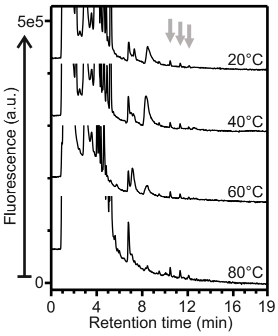


**Negative control - K49**

**Capsule type:** group 4
Wzx/Wzy-dependent
**Strain**: *E. coli* reference strain K37
**Strain No.**: SSI #81995
**Serotype**: O8:K49:H21
**Culture media**: LB
**Approx. per injection**: 2e8 cells

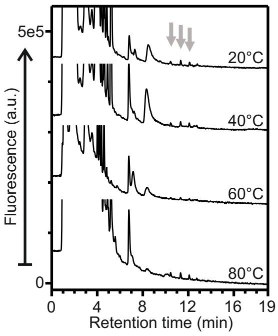


**Negative control - K50**

**Capsule type:** group 4
Wzx/Wzy-dependent
**Strain**: *E. coli* reference strain K50
**Strain No.**: SSI #81996
**Serotype**: O8:K50:H-
**Culture media**: LB
**Approx. per injection**: 2e8 cells

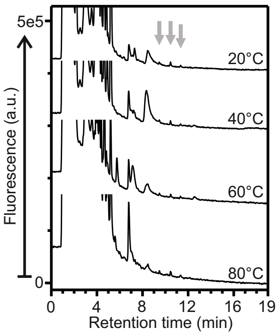


**Negative control - K55**

**Capsule type:** group 1
Wzx/Wzy-dependent
**Strain**: *E. coli* reference strain K55
**Strain No.**: SSI #82001
**Serotype**: O9:K55:H-
**Culture media**: LB
**Approx. per injection**: 2e8 cells

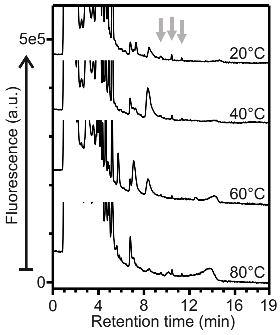


**Negative control - K83**

**Capsule type:** group 4
Wzx/Wzy-dependent
**Strain**: *E. coli* reference strain K83
**Strain No.**: SSI #82003
**Serotype**: O20:K83:H26
**Culture media**: LB
**Approx. per injection**: 2e8 cells

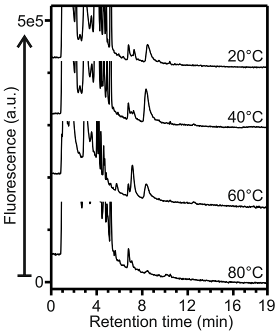


**Negative control - K87**

**Capsule type:** group 4
Wzx/Wzy-dependent
**Strain**: *E. coli* reference strain K87
**Strain No.**: SSI #82005
**Serotype**: O8:K87:H19
**Culture media**: LB
**Approx. per injection**: 2e8 cells

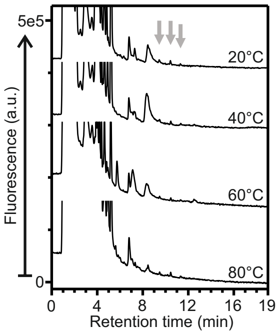


**Negative control - K101**

**Capsule type:** group 4
Wzx/Wzy-dependent
**Strain**: *E. coli* reference strain K101
**Strain No.**: SSI #82008
**Serotype**: O20:K101:H-:F6
**Culture media**: LB
**Approx. per injection**: 2e8 cells

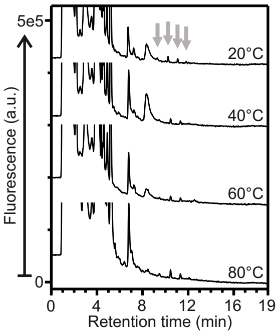


**Negative control - K102**

**Capsule type:** group 1
Wzx/Wzy-dependent
**Strain**: *E. coli* reference strain K102
**Strain No.**: SSI #82009
**Serotype**: O8:K102:H-
**Culture media**: LB
**Approx. per injection**: 2e8 cells

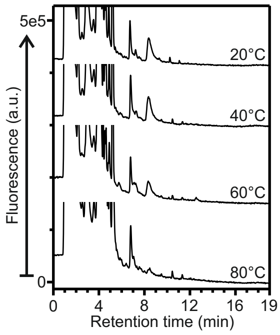


**Negative control - K103**

**Capsule type:** group 1
Wzx/Wzy-dependent
**Strain**: *E. coli* reference strain K103
**Strain No.**: SSI #82010
**Serotype**: O101:K103:H-
**Culture media**: LB
**Approx. per injection**: 2e8 cells

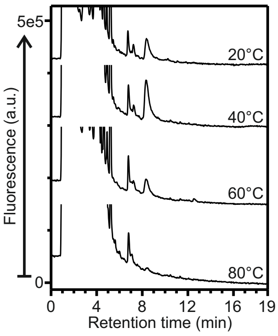
